# Adeno-associated Virus Receptor-binding: Flexible Domains and Alternative Conformations through cryo-Electron Tomography of AAV2 and AAV5 complexes

**DOI:** 10.1101/2022.01.10.475736

**Authors:** Guiqing Hu, Mark A. Silveria, Michael S. Chapman, Scott M. Stagg

**Affiliations:** Institute of Molecular Biophysics, Florida State University, FL 32306, USA; Department of Biochemistry, University of Missouri, Columbia, MO 65211, USA; Department of Biological Sciences, Florida State University, FL 32306, USA

## Abstract

Recombinant forms of adeno-associated virus (rAAV) are vectors of choice in the development of treatments for a number of genetic dispositions. Greater understanding of AAV’s molecular virology is needed to underpin needed improvements in efficiency and specificity. Recent advances have included identification of a near universal entry receptor, AAVR, and structures by cryo-electron microscopy (EM) single particle analysis (SPA) that revealed, at high resolution, only the domains of AAVR most tightly bound to AAV. Here, cryogenic electron tomography (cryo-ET) is applied to reveal the neighboring domains of the flexible receptor. For AAV5, where the PKD1 domain is bound strongly, PKD2 is seen in three configurations extending away from the virus. AAV2 binds tightly to the PKD2 domain at a distinct site, and cryo-ET now reveals four configurations of PKD1, all different from that seen in AAV5. The AAV2 receptor complex also shows unmodeled features on the inner surface that appear to be an equilibrium alternate configuration. Other AAV structures start near the 5-fold axis, but now β-strand A is the minor conformer and, for the major conformer, partially ordered N-termini near the 2-fold axis join the canonical capsid jellyroll fold at the βA-βB turn. The addition of cryo-ET is revealing unappreciated complexity that is likely relevant to viral entry and to the development of improved gene therapy vectors.

**IMPORTANCE:** With 150 clinical trials for 30 diseases underway, AAV is a leading gene therapy vector. Immunotoxicity at high doses used to overcome inefficient transduction, has occasionally proven fatal and highlighted gaps in fundamental virology. AAV enters cells, interacting through distinct sites with different domains of the AAVR receptor, according to AAV clade. Single domains are resolved in structures by cryogenic electron microscopy. Here, the adjoining domains are revealed by cryo-electron tomography of AAV2 and AAV5 complexes. They are in flexible configurations interacting minimally with AAV, despite measurable dependence of AAV2 transduction on both domains.

## Introduction

Adeno-Associated Virus (AAV) is a small 25nm T=1 icosahedral virus with a protein shell encapsidating a single-stranded DNA genome (1, 2). Adeno-Associated Virus is so named as it was discovered during adenovirus preparations and its replication depends on co-infection with adenovirus or one of several other “helper” viruses, not because there is any structural relation (3-5). AAVs were long regarded as non-pathogenic, an initial rationale for their development as transducing vectors for *in vivo* (and *ex vivo*) gene therapy. (6-9). Recent discovery of AAV sequences inserted into proto-oncogenes of patients with hepatocellular carcinoma (HCC) has prompted vigorous debate about causal links to natural infection and future vector use (10-16). A prevalent view is emerging that there may be a concern for individuals with chronic liver disease (17-19).

Nonetheless, it is exciting times for gene therapy. After many years in development, the first two *in vivo* treatments have been approved by the U.S. Food and Drug Administration (FDA), using AAV2 and AAV9 vectors respectively. Luxterna™ is a treatment for an inherited blindness, Zolgensma™ for spinal muscular atrophy (20, 21). AAV vectors are being used for > 150 ongoing clinical trials (https://clinicaltrials.gov/) (22), but challenges await in generalization of the early successes. Deaths in a myotubular myopathy trial likely resulted from immune-toxicity of the high doses needed to achieve therapeutic expression levels with an inefficient transducing vector (23, 24). Structural studies are key to an improved fundamental understanding of AAV’s virology, and in its engineering for vector improvement.

Initial crystallographic structures revealed the 60-fold symmetric part of the capsid. The capsid gene is expressed as three variant viral proteins (VP) due to alternative start codons and splice variants (25). The variants are in-frame, sharing most of their amino acid sequences, and it has become conventional to use common numbering, based on the largest, VP1. Ordered structure becomes visible at about residue 220, or ∼20 residues beyond the N-terminus of VP3 that constitutes ∼80% of the capsid (26). Upstream, VP1 (∼10%) and VP2 (∼10%) are extended by a common region of 65 usually unseen amino acids that some have proposed to function in nuclear localization (27, 28). Then there is a segment unique to VP1 (VP1u), N-terminal of the VP2 start, that contains a phospholipase A2 (PLA_2_) domain that is initially sequestered within the capsid, but becomes exposed for endosomal escape on the entry pathway (29-33).

Over 130 variants of human and non-human primate AAVs have been identified (34, 35). These are grouped into eight major named and unnamed clades, containing one or more serotypes that are antigenically distinct, i.e. antibodies recognizing one serotype do not cross-react with others (34, 36). The serotypes differ in other properties, such as binding preference to glycan attachment factors and empirically determined tissue tropisms (37, 38). This study uses two representatives, AAV2 and AAV5 as model systems. AAV2 is the type species that is the best characterized. AAV5 is tied for the most distantly related with reference to VP3 amino acid sequence.

This study further characterizes interactions with the near-universal protein receptor, AAVR. AAVR was only recently discovered though unbiassed genome-wide screening as a receptor key for entry and trafficking (39). Previously, a serotype-specific variety of glycans had been considered to be “primary receptors”, heparan sulfate proteoglycan (HSPG) for AAV2 and sialic acid (SIA) for AAV5 (40-42). However, it has recently been argued that the glycans have less specific roles than classic receptors, and, following virological convention, should be considered attachment factors (43), anchoring viruses to cell surfaces but not mediating entry. Several membrane proteins, primarily tyrosine kinase receptors and integrins, were also identified as co-receptors for different serotypes, but they have not figured in several more recent knock-out screens (39, 44-53). Current evidence indicates that AAV2 and AAV5 attach to cells using different extracellular glycans, that both depend on AAVR for entry and trafficking, and that then AAV2 (but not AAV5) has a downstream dependence on another host membrane protein, GPR108 (53, 54).

AAVR is a C-terminally anchored transmembrane protein, in which the ectodomain (from the N-terminus) consists of a signal peptide, a MANEC domain (motif at N-terminus with eight cysteines) then five Ig-like PKD (polycystic kidney disease) domains (55, 56). It is the PKD domains that bind AAV, but surprisingly there are different serotype-specific domain dependencies (54). For AAV2, PKD2 is most important, but PKD1 has an accessory role, whereas AAV5 is exclusively dependent upon PKD1 (54). These determinations were made by: (a) SPR measurements using AAV and heterologously expressed AAVR domain fragments; (b) transduction inhibition through addition of solubilized domain fragments; (c) knock-out through domain-deletion; and (d) viral overlay assay (39) Concurrent cryogenic-Electron Microscopy (cryo-EM) structure determinations using different expressed AAVR fragments, PKD1-5 or PKD1-2, revealed PKD2, bound to AAV2 at 2.8 and 2.4 Å respectively (43, 57). Even though the samples contained 5- and 2-domain fragments respectively, only the most tightly interacting domain (PKD2) was revealed. Cryo-electron tomography (cryo-ET) of an N-terminal fusion of maltose-binding protein (MBP) and PKD1-5, combined with cross-linking mass spectrometry (XL-MS) was consistent, showing anchoring of PKD2 to the viral surface, and the PKD3-5 domains emanating radially in at least four configurations (43). Then, in succession, came cryo-EM structures of AAV5 complexes, PKD1-5 at 3.2 Å and PKD12 at 2.5 Å resolution, now showing just the PKD1 domain that alone had previously been implicated (54, 58, 59). Intriguingly, the homologous PKD1 and PKD2 domains were not accommodated as variations of a single AAVR binding site on AAV, but at distinct sites. One could then best imagine evolutionary divergence occurring through an ancestral form that bound both domains, but overlay of the structures eliminated simple explanations with the finding that the domains could not be connected plausibly by the unseen five-residue linker (59).

Cryo-ET has technical advantages enabling determination of 3D structures of flexible molecules in heterogeneous configurations, such as AAVR with its variable PKD domain orientations. In contrast to single particle cryo-EM, where a single 2D image from many identical or nearly identical particles (10^4^ to 10^6^) are aligned and averaged into a 3D reconstruction, in tomography, 3D images of every individual particle are realized by tilting the microscope stage. This technique has some limitations because the sample can only be tilted within a range of angles between -65° and +65°. A consequence of this is that the resulting 3D reconstructions have a “missing wedge” of information that can distort the 3D volumes. However, the missing wedge can be filled by averaging between aligned subvolumes containing a structure of interest in different orientations and thus with different missing wedges. A structure can be split into subvolume parts for classification and averaging to characterize variability in heterogeneous regions. This can be a particular advantage for structures like virus-receptor complexes where different copies of a viral capsid protein could have receptor bound in a different configuration. There have been several successful applications of the approach, to, for example, the heterogenous structure of Simian Immunodeficiency Virus (SIV) envelope glycoprotein when bound by CD4 receptor or monoclonal antibody 36D5 (60).

Here we use cryo-ET to focus on the 2-domain receptor complex of AAV for a holistic and hybrid comparison with single particle cryo-EM to locate the parts that had been refractory to the high resolution cryo-EM. It is using the unique advantages of cryo-ET to distinguish different conformational states, focusing reconstructions on the sub-volumes surrounding each 3-fold axis to reveal the hitherto unseen domains in the AAV-PKD12 complexes, and other elements of both the receptor and virus structures that have been smeared beyond recognition in the 60-fold averaged cryo-EM reconstructions.

## Materials and methods

### Virus and receptor preparation

Virus like particles (VLPs) for both AAV2 and AAV5 were prepared as previously described (43, 59, 61). In brief, VLPs, which consist of the protein shells absent the viral DNA, were expressed in Sf9 cells using the Invitrogen Bac-to-Bac protocol and a pFastBacLIC cloning vector (addgene #30111). VLPs then underwent three rounds of CsCl density gradient ultracentrifugation. An additional step of heparin affinity chromatography was performed on the AAV2 VLPs.

AAVR constructs were expressed using a pET-11a vector in E. coli BL21(DE3) cells as described previously (43, 61). PKD1-2 was expressed with a 6x histidine tag and purified using Co^2+^ affinity chromatography.

### Cryo-electron microscopy

Complexes of AAV2 with AAVR PKD1-2 were prepared on-grid as follows. A thin layer of carbon was deposited on a mica sheet using a Cressington Carbon Coater and was floated onto Quantifoil R1.2/1.3 grids. Then the carbon-coated Quantifoil grids were glow-discharged for 20 s with a Solarus 950 (Gatan). 4 μL of AAV2 VLP at concentration 0.2 mg/ml was applied on grid and incubated for 1.5 minutes. The grid was gently blotted on the side with filter paper, and another 4 μL of PKD1-2 at concentration 1.5 mg/ml was applied and allowed to incubate for another 1.5 minutes followed by plunge-freeze using a Vitrobot Mark IV (FEI).

EM grids of AAV5 complexed with AAVR PKD1-2 were prepared in a similar way. VLP and receptor were first dialyzed into 25 mM HEPES, 150 mM NaCl, pH 7.4. EM samples were then prepared on glow discharged ultrathin continuous carbon film supported by lacey carbon on copper grids (Ted Pella, Redding, CA, USA, Cat No 01824). First 2 μL of ∼5.4 μM AAV5 VLP was added to the grid and given 2 minutes to adhere. Sample was then wicked and 2 μL of 33 μM PKD1-2 was added. Grids were then plunge frozen using an FEI Vitrobot Mark IV with a blot force of 4, time of 2 sec, temperature of 25 °C, and at 100% humidity.

Cryo-ET tilt series were acquired on an ThermoFisher Titan Krios (Hillsboro, OR) and recorded with Leginon software (Suloway et al., 2009) on a Gatan K3 direct detector. A magnification of 33,000 X was used with a pixel size of 2.74 Å and a total dose of 100 e^—^/Å² per tilt series. The tilt angle ranged from -60° to 60° with 2° steps. Exposure time at each tilt step was automatically adapted by the Leginon software according to the tilt angle. The number of frames at each tilt step was automatically set by Leginon according to the exposure time at each tilt step. Dose was fractionated across the frames at each step. Defocus values were set to 5 μm underfocus.

Single particle data was also collected on the Titan Krios with the K3 camera using Leginon software. Magnification was 81,000 X and pixelsize was 1.1 Å. The defocus range was set to -1.0 to -3.0 μm. Total dose was ∼60 e^—^/Å² per image with 50 frames for each micrograph. All frames of each micrograph were aligned using MotionCor2 (62).

For AAV5 bound with PKD1-2, both single particle and tomography data were collected from the same cryo-EM grid.

### Single particle image processing

CTFFIND4 and GCTF were used to estimate contrast transfer function (CTF) parameters on all motion-corrected micrographs and the best estimate was chosen using resolution evaluation in Appion (63-65). Around 1000 particles were picked using DoG (Difference of Gaussian) Picker and the rotational average of those particles was used as the template for picking using FindEM in Appion (65, 66). A total of 26,091 particles were picked from 378 micrographs and extracted with a box size of 432 × 432 pixels in Appion. 2D classification and 3D classification were conducted to choose good particles in Relion3-beta. The previous 2.5 Å resolution single particle cryo-EM reconstruction of the AAV5-PKD12 complex (EMD22988) was low pass filtered to 60 Å resolution and used as the initial reference for 3D refinement by Relion3-beta (43, 67). A total of 15,052 particles were selected for gold standard auto-refinement. Icosahedral symmetry was applied for auto-refinement. After auto-refinement, CTF refinement and beam tilt refinement, a final map of 2.88 Å resolution was achieved.

### Tomography image processing

Tilt series were aligned using Protomo software within Appion (Winkler and Taylor, 2006; Lander et al., 2009; Noble and Stagg, 2015). Following that, the image stack for tilt series were imported into EMAN2/e2tomo. Alignment parameters from Protomo were imported into EMAN2/e2tomo with home-made scripts, and tomograms were directly calculated with imported parameters. Contrast transfer function (CTF) parameters were estimated on all micrographs inside E2tomo. For AAV2 bound with PKD1-2, 127 virions (7620 asymmetric subunits) were manually picked and extracted with E2tomo and a box size 288 × 288 pixels. Similarly, 85 virions (5100 asymmetric subunits) were picked and extracted for AAV5 bound with PKD1-2. The same 2.4 Å AAV2-PKD12 reconstruction (EMD-0553) was low pass filtered, now to 50 Å and used as the initial reference for alignment. For the complex of PKD1-2 with AAV5, extracted 3D subtomograms were aligned using the new 2.88 Å SPA reconstruction of the PKD12-AAV5 complex, low pass filtered to 50 Å. Subtilt refinement was then used to align the individual 2D particle images in each tilt, and apply a per-particle-per-tilt CTF correction. The sub-tilt refined 3D particles were exported from EMAN2/e2tomo and then imported into the program I3 (68, 69). Icosahedral symmetry was applied to generate 60 copies of each particle such that each possible asymmetric unit was overlaid onto the same frame of reference. The particles were then translated and rotated to center on the 3-fold spike in a “spike up” standard orientation. At this point, particles were re-extracted with box size of 90 × 90 pixels surrounding a single 3-fold spike, facilitating classification of asymmetric units. In order to improve the signal/noise ratio, the re-extracted subtomograms, containing one trimer, were binned by 2. Then classification was conducted on one asymmetric unit to reveal the PKD1 domain (AAV2 bound with PKD1-2) or PKD2 domain (AAV5 bound with PKD1-2). Even though the resulting classes are based on a single asymmetric unit, the classes were re-expanded by three-fold symmetry to better illustrate the context of the extra domains.

### Model fitting

All the tomography maps for AAV2 and AAV5 bound with PKD1-2 were aligned to the same frame of reference. High resolution SPA reconstructions EMD0553 and EMD9672 were aligned with the global (overall) average of the subtomograms of AAV2 bound with PKD1-2 (43, 57). Similarly, the map for the newly obtained 2.88 Å single particle reconstruction for the AAV5/PKD1-2 complex was aligned to the global average subtomogram of the same complex. PDBid 6ihb, with its well-ordered βA, was used for the atomic model of the AAV2 viral protein, while PKD2 was taken from the higher resolution PDBid 6nz0 (43, 57). Likewise, the atomic model of VP protein and PKD1 from PDBid 7kpn was docked as a single rigid body trimer into the newly obtained 2.88 Å reconstruction for the AAV5/PKD1-2 complex and used to interpret the tomography maps. In summary, the tomographic reconstructions for AAV2 and AAV5 complexes were calculated in the same frame of reference, then high resolution SPA reconstructions and atomic models were overlaid, facilitating comparisons.

The previously unseen domains were modeled using the following. Atomic models for PKD1 and PKD2 were taken from PDB entries 7kpn and 6nz0 respectively, and were fitted, as rigid domains, into the newly revealed domain densities separately for the 4 AAV2 classes and the 3 AAV5 classes using Chimera (70).

### Contact analysis

The VMD “atomselect” command was used to identify additional potential residue contacts contributed by the AAVR domains that had not previously been resolved in single particle analysis (Humphrey et al., 1996). Distances between the newly revealed PKD1 (AAV2) / PKD2 (AAV5) and respective viral proteins were calculated. The “atomselect” command lists the residue numbers of all residues that have any atom approaching within 4.5 Å.

## Results

### The structure of AAV2 bound with PKD1-2

Cryogenic electron tomographic (cryo-ET) tilt series were acquired for AAV2 bound by the PKD1-2 domains of AAVR and tomograms were reconstructed. AAV2 subvolumes were aligned and then subdivided into individual trimers and all aligned trimers were averaged (Figure 1B_1_). This revealed densities at about 20 Å resolution, corresponding to the VP capsid protein and PKD2, consistent with the published single particle reconstructions of the AAV2/PKD1-2 complexes (43, 57). The atomic model of the AAV2 spike fits the global average well, defining the density for the viral protein and the PKD2 domain of AAVR (Figure 1B_2_). As with the single particle analysis, the AAV2 viral protein and the PKD2 domain are readily apparent, but there was no sign of the PKD1 domain. In order to reveal PKD1, tomographic subclassification was performed using a trimer subvolume that extended mostly outwards beyond PKD2 for the AAV2/PKD1-2 complex. This revealed additional features corresponding to four distinct conformations for PKD1 (Figure 1C). The features are of the correct shape and size for a PKD domain, and an atomic model of PKD1 fits well into the map of each class (Figure 1D). A 5 residue linker (residues 400-404) can be built between the PKD1 and PKD2 domains with plausible stereochemistry.

**Figure 1:**
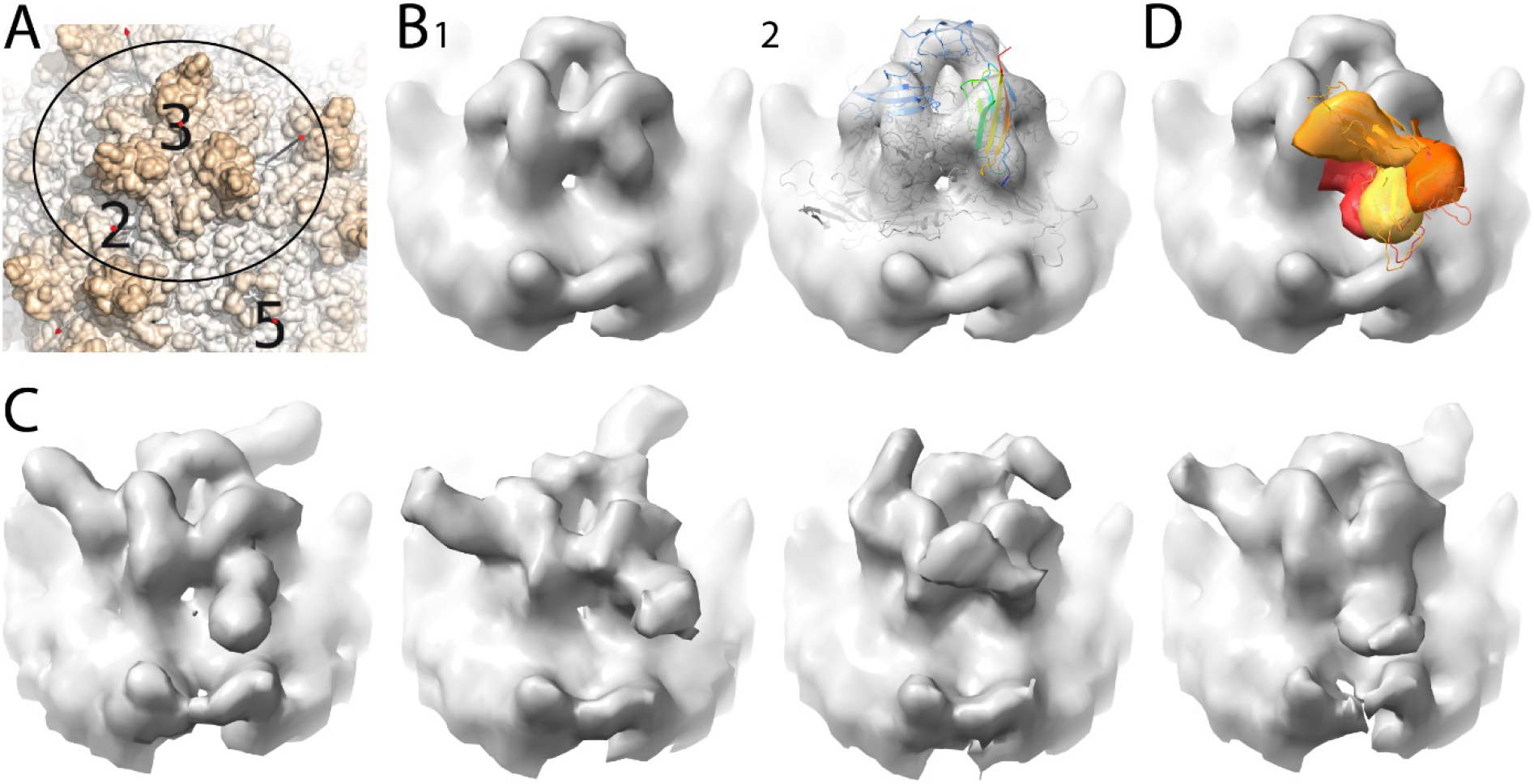
Classification of AAV2/PKD1-2 trimer spikes. A) Surface representation of AAV2. The circle shows the area that was classified by subvolume averaging and the axes of symmetry are labeled as 2, 3, and 5. B) Global tomographic subvolume average of AAV2/PKD1-2 trimeric spike as solid surface (B1) and translucently (B2) overlaid on the atomic model of a subunit trimer from the prior single particle reconstruction (PDB ID 6nz0). The viral proteins are shown in grey ribbons and two of the PKD2 domains are colored blue, while one is rainbow colored by residue number from N (blue) to C (red). C) Classification of the region outside of the PKD2 domain revealed 4 distinct conformations for PKD1. D) The 4 PKD1 domain conformations were segmented and overlaid and PKD1 atomic models (shades of orange) fit to the classes seen in C1-C4.

### The structure of AAV5 bound with PKD1-2

Cryo-ET, subvolume averaging, and classification were performed on AAV5 in complex with the PKD1-2 domains similarly to AAV2/PDK1-2. The global average of aligned subtomograms of AAV5 bound with PKD1-2 showed clear density for the PKD1 domain, in agreement with the previously published single particle cryo-EM reconstructions of AAV5/PKD1-2 (58, 59). Fitting the viral trimer spike model and PKD1 into this averaged tomographic map revealed features that were unaccounted by the atomic models for the viral protein and PKD1 domain. These features were in positions extending away from the surface of the capsid that were plausible locations of PKD2. This differs from the AAV2/PDK1-2 tomography, where density for the “missing” domain (PKD1 for AAV2) was only revealed on classification and was not apparent in the global average. This indicates that the PKD2 domain is more constrained when bound to AAV5 than PKD1 is when it is bound to AAV2. The PKD2 domain becomes better defined upon classification, focusing on the area outside PKD1, which yielded three distinct classes (Figure 2B). As with AAV2, the extra densities are the correct shape and size for a PKD domain, and an atomic model of PKD2 fits well into the class maps (Figure 2C). In each case, the first residue of the PKD2 model is within 19 Å of the C-terminal residue of PKD1, close enough to be bridged by the 5-residue domain linker. Variation in PKD2 orientation among classes of the AAV5 complex is modest (Figure 2C) which is consistent with conformations that are more constrained than the AAVR conformations that we observed with AAV2.

**Figure 2:**
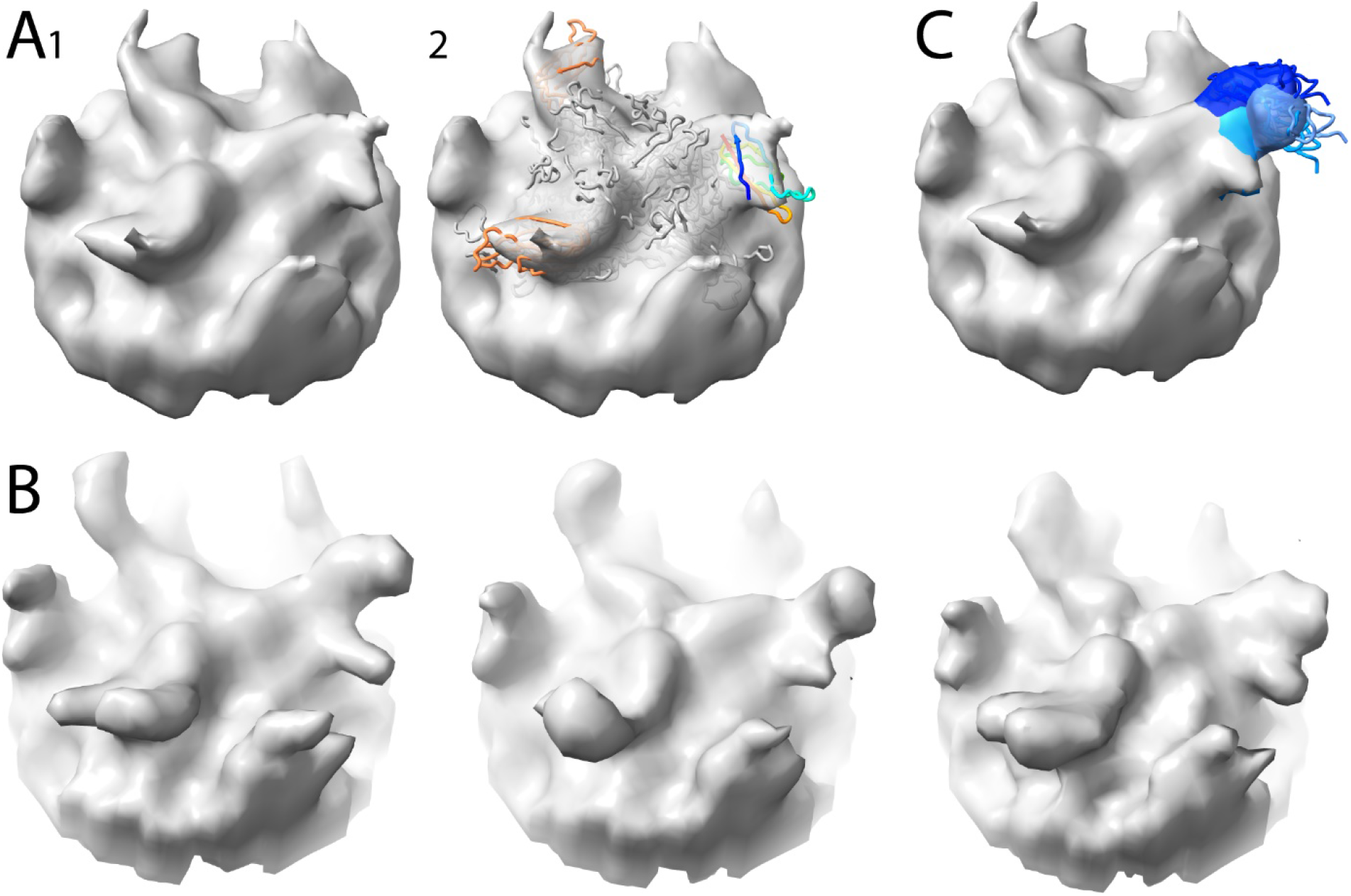
Classification of AAV5/PKD1-2 trimer spikes, oriented as in Figure 1A. A) Global tomographic subvolume average of the AAV5/PKD1-2 trimeric spike, alone (A1) and overlaid (A2) with the atomic structure from single particle reconstruction (PDBid 7kp3) in which the PKD1 domain of AAVR was seen (orange). B) Classification of the region outside of the PKD1 domain revealed 3 distinct conformations for PKD2. C) The 3 PKD2 domain conformations were segmented and shown overlapping (shades of blue).

### Hybrid analysis: Integration with Cross-linking Mass Spectrometry (XL-MS)

Meyer et al. reported mass spectroscopic identification of amino acids in AAV2 and AAV-DJ that were cross-linked with cyanurbiotindimercaptopropionylsuccinimide (CBDPS) that has a spacer length of 14 Å (43). Atomic models can be compared with these distance constraints, with the caveats that the tomographic classes are at ∼20 Å resolution and without definition of side chains, and that the distances are measured from cryo-ET samples that were not cross-linked, so do not reflect any remodeling of local structure on cross-linking (Table 1).

**Table 1:** Consistency of the atomic models built into the AAV2-AAVR cryo-ET with cross-linking mass spectrometry (XL-MS). Under each class is listed the observed distance between the reactive groups on AAVR and AAV2.

| XL-MS experiment |  |  |  |  | Distance measured from structure |  |  |  |  |
| --- | --- | --- | --- | --- | --- | --- | --- | --- | --- |
| Virus | Virus residue | AAVR construct | Receptor residue | Location | Class 1 | Class 2 | Class 3 | Class 4 | Measured from: |
| AAVDJ | K558<br>(equivalent to AAV2 K556) | Full ecto-protein | K404 | PKD2 (N-terminal) | 13 Å | 13 Å | 13 Å | 13 Å | PKD2 modeled into high resolution EM of AAV2-PKD1/2 |
| AAV2 | K490 | MBP-PKD1-5 | K399 | PKD1 | 37 Å | 30 Å | 26 Å | 15 Å | PKD1 docked into cryo-ET |
| AAV2 | T560 | MBP-PKD1-5 | 399 | PKD1 | 43 Å | 37 Å | 40 Å | 23 Å | As above |
| AAV2 | K556 | MBP-PKD1-5 | K338 | PKD1 | 61 Å | 72 Å | 36 Å | 59 Å | As above |
| AAV2 | T450 | MBP-PKD1-5 | K399 | PKD1 | 38 Å | 38 Å | 33 Å | 24 Å | As above |

Given that at best nanometer-level consistency should be expected, class 4 provides plausible explanation for 4 of 5 observed cross-links, one involving the tightly bound PKD2 and three involving the C-terminus of PKD1 which is closest to PKD2 and the virus surface. A rationalization of K338 cross-linking (minimal 36 Å) is more tenuous, requiring remodeling of the lysine side chains. It seems more likely that tomography is sampling four of many possible PKD1 orientations, and that any of the larger population could be captured in cross-linking. In other words, the cross-linking reflects a highly flexible receptor with many domain orientations, of which a subset, perhaps the most stable, are sampled in the tomographic classes.

### Comparison of the AAV2 and AAV5 complexes with PKD1-2

When AAV2 is bound by PKD1-2, the PKD2 domain has the highest affinity but PKD1 has measurable impact, while, for AAV5, PKD1 appears to be the only domain involved (54). These results, coming from binding and transduction analysis of domain-swap and deletion mutants are supported and rationalized by the current tomography study. The tomography reveals other differences in the PKD1/PKD2 domain modes of binding to the two serotypes. For PKD1-2 bound to AAV2, no density is revealed for the PKD1 domain in the global average of aligned subtomograms, indicating a high level of heterogeneity of the PKD1 domain. For the AAV5 complex, even though weak, density of the previously-missing PKD2 domain is apparent in the global average of the aligned subtomograms. This difference between global averages indicates that PKD1 in AAV2 is more heterogeneous than PKD2 in AAV5. This is further confirmed by the classes for the extra PKD1/PKD2. As shown in Figure 3, the extent of variability in AAV2 is much higher than AAV5 with a wider range of orientations. Furthermore, for PKD1-2 bound to AAV2, three out of four of the classes are in extended conformations with obtuse angles between the two PKD domains and the fourth class has the two PKD domains folded back on each other. For PKD1-2 bound to AAV5, the two domains are always at an acute angle, folded back towards one another and contacting near the hinge in an antiparallel hairpin configuration. The extra PKD1/PKD2 also differs on the contact with viral proteins. Consistent with the accessory role of PKD1 in AAV2 cellular entry, one the four classes of PKD1 appears to have some contact with the viral protein, whereas for AAV5 bound there appear to be no contacts with PKD2 (see below).

**Figure 3:**
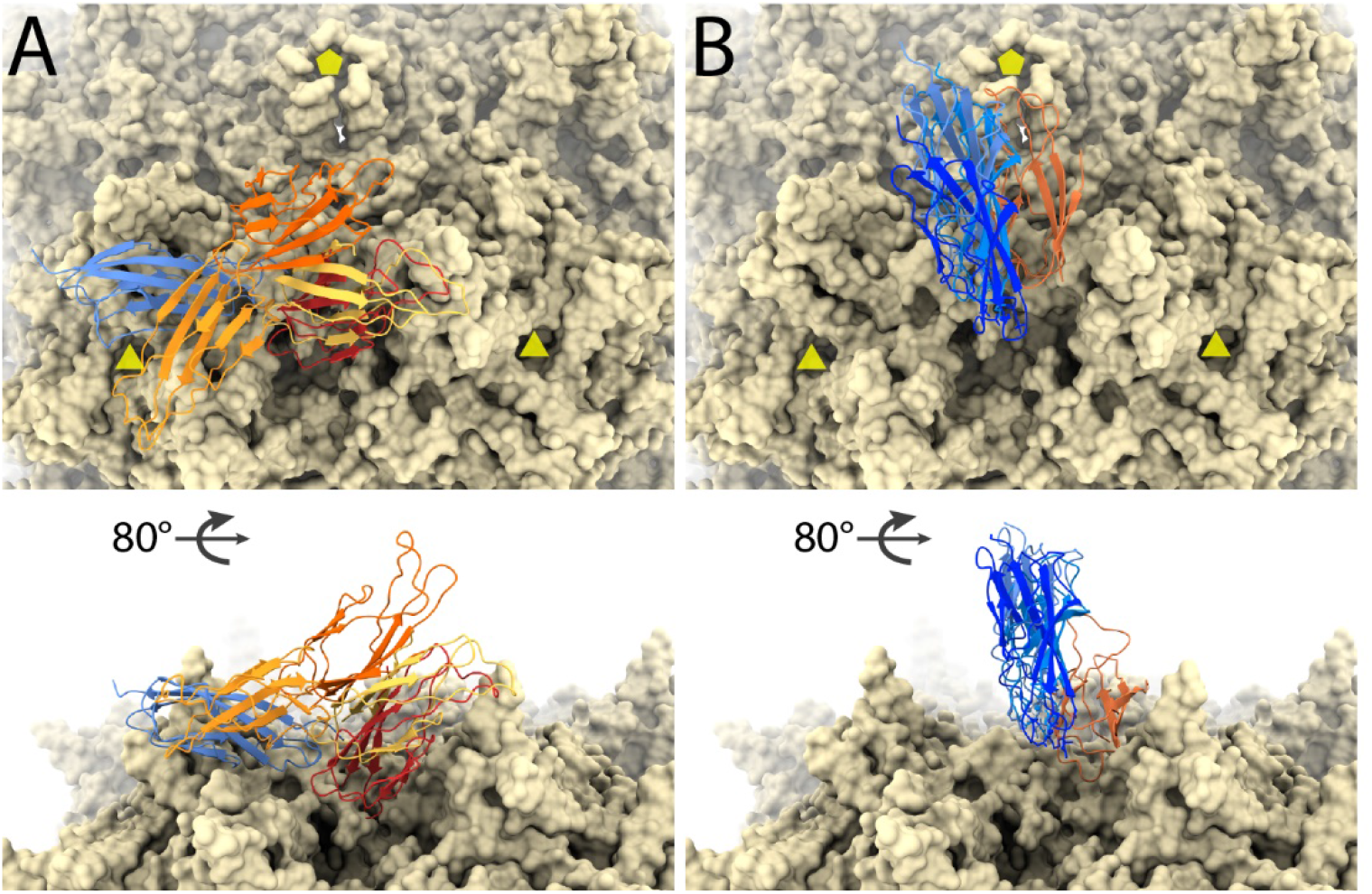
Conformational variability of AAV2/PDK1-2 and AAV5/PKD1-2 structures. The virus surfaces are viewed along a 2-fold axis (top) and tangentially (bottom), with 3-folds and 5-folds marked as triangles and pentagons. In each, the AAVR receptor is shown bound to one of the symmetry-equivalent regions on the virus surface. A) AAV2, with the AAVR PKD2 domain (blue) tightly bound and the preceding PDK1 domain in several orientations (shades of orange) as determined from the subvolume tomography. B) In its complex with AAV5, a tightly bound conformation of AAVR PKD1 (orange) is followed by PDK2 in three orientations (shades of blue) that are more constrained compared to PKD1 in (A).

### Contact between newly revealed extra PKD1/2 with VP protein

The details of interactions between PKD2 with AAV2 and PKD1 with AAV5 have been discussed in the respective high resolution single particle cryo-EM analyses (43, 59). Here, contact analyses are added for the newly revealed flexible PKD1 in its AAV2 complex and PKD2 in its AAV5 complex, but with a caveat that needs to be emphasized. The resolution of the tomographic classes (and therefore precision of atomic models) is low, ∼20 Å. One must therefore be very cautious in interpreting whether 4.5 Å contact distance criteria are met for individual pairs of amino acids and the analysis for each class is more appropriately an indicator of whether contacts are extensive or minimal. For the AAV2 class 4 model, only three residues of PKD1 (R353, V398 & K399) approach AAV2 closely (near E385 & D529 from different subunits). For the other three classes of AAV2 complex, there appear to be no contacts with PKD1 beyond the contacts already established at high resolution for the tightly bound PKD2 domain (43). These results are consistent with the cross-linking data in Table 1. For the classes of the AAV5 complex, no contacts are apparent between PKD2 and the viral protein.

### Features on the inside surface of AAV2

In the global average of the aligned subtomograms of AAV2 bound with PKD1-2, there is extra density that projects in toward the center of the virus that is not accounted for by the known AAV atomic models (Figure 4). Interestingly, the extra density was only observed in our AAV2/PKD1-2 structure (Figure 4A) but not the AAV5/PKD1-2 structure (Figure 4B). Model fitting shows that this protruding density is located in close proximity to residue 237 of the VP protein (Figure 4C). Maps for two previous AAV2/PKD1-2 single particle cryo-EM reconstructions are mostly quite similar, but differ here. The Zhang et al. structure at 2.4 Å (EMD9672) is similar to most prior AAV structures, interpretable from residue 219, with β-strand A, running anti-parallel to βB before a hairpin turn at Gly_236_-Asp_237_ that connects the two strands (57). This map for βA is slightly weaker than βB and other strands, but only slightly, with only a slight hint of disorder (Fig. 4D). By contrast, in the 2.4 Å structure of Meyer et al. (EMD0553), there is little in the map to suggest the presence of βA (Figure 4C) (43). A small fraction of subunits have a βA configured as in the Zhang et al. structure, as evidenced by clear, but weaker features for the distinctive Trp_234_. However, N-terminally of Trp_234_, little is recognizable, and, in deference to the map weakness indicating low order and/or low occupancy for the βA configuration, βA is not included in the atomic model.

**Figure 4:**
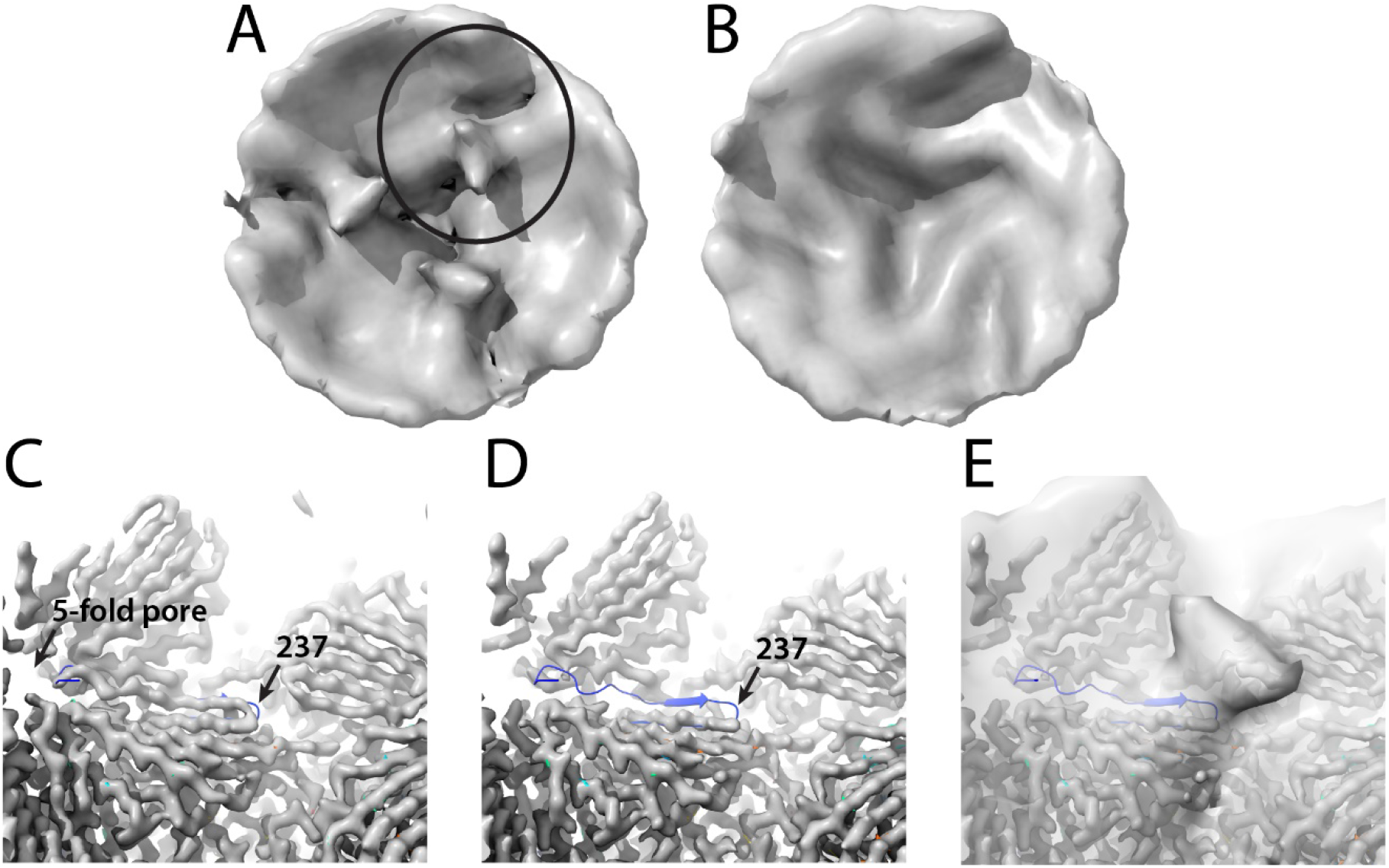
Unmodeled density inside the AAV2 capsid. A) The inner surface of a subvolume average of the AAV2/PKD1-2 complex surrounding a 3-fold, viewed outwards. The circled region is highlighted in C-E. B) The corresponding subvolume of the AAV5/PKD1-2 complex. C) The 2.8 Å single particle reconstruction (EMD9672) and model (PDBid 6ihb) of an AAV2-AAVR (PKD1-5) complex (57). The map clearly shows βA proceeding left-to-right to the hairpin turn at residue 237 as has been seen in AAV2 crystal structures (1). D) In the 2.4 Å single particle reconstruction of a AAV2-AAVR complex (PKD1-2; EMD0553) (43), this configuration of βA is not apparent, as illustrated by overlaying the (mismatched) model PDBid 6ihb (24, 57). E) Overlaid on panel D is now the newly observed AAV2/PKD1-2 subvolume average tomographic density, suggesting that the unmodeled density extending inwards from the surface is a partially ordered configuration of upstream residues. It would be an alternate to the βA strand, connecting in to βB at residue 237.

Relevant to these observations, it has been established that the AAV capsid is assembled from VP1, 2, and 3 in a roughly 1:1:10 ratio, sharing much of their sequence and structure, but differing in their N-terminal extensions (see Introduction) (71). Disordered features in the same general area were first seen as “fuzzy globules” in a 2001 nanometer-resolution SPA reconstruction of empty AAV2 virus-like particles (72). Assignment as parts of VP1 and/or VP2 was supported by structures lacking “fuzzy globules” either for mutants in which VP1/2 were deleted, or in capsids following heat treatment that was known to expose VP1u on the exterior of AAV (73). However, doubts emerged with the absence of the “fuzzy globules” in reconstructions of AAV1 vectors with varying DNA content, again by the same group (74). For the most part, these disordered features have not been noted in subsequent structures. However, they did resurface in the cryo-EM reconstruction of the AAV2 R432A mutant, in which map was missing at 3.7 Å resolution for βA. When this map was viewed at 5 Å resolution, there was feature extending from the 236-7 hairpin turn that was interpreted as four residues extending toward the general area of the “fuzzy globules” (75). The map at 5 Å resolution had not been deposited, so for comparison to our cryo-ET, the 3.7 Å map (EMD8100) was low-pass filtered to 11 Å resolution. At this lower resolution, we see not just 4 amino acids heading from the βA-βB turn towards the 2-fold (75), but additionally we see the larger unmodeled feature that is also present in the AAV2-PKD12 tomography. Furthermore, this feature is the same as the “fuzzy globules” seen earlier in AAV2 VLPs at 11 Å resolution (72). For both our PKD12 complex and the R432A packaging mutant, higher resolution cryo-EM SPA indicates that the presence of the unmodeled feature is accompanied by loss of βA, i.e. it is an alternative configuration for the N-terminal residues. In the prior 3.8 Å reconstruction of the wild-type AAV2, βA was clear, and we now add that when low-pass filtered to 11 Å, EMD8099 shows is no indication of the partially ordered alternative configuration seen in the R432A mutant (75).

Whereas we see the partially ordered alternative configuration in our complex of AAV2 with the PKD1-2 fragment of AAVR, the nominally similar complex of AAV2 with a PKD1-5 fragment has density for βA that is only marginally weaker than βB (EMD 9671) (57), so the more usual configuration of βA predominates. Thus, there is not a simple and deterministic receptor-triggered conformational switch. Indeed weak, but recognizable density for Trp_234_ in the 2.4 Å structure of the PKD1-2 complex (43) indicates an equilibrium (favoring the alternative configuration) that might reflect incomplete PKD1-2 binding or an intrinsic and perhaps dynamic finely-balanced equilibrium. The latter is consistent with a history of VLP structures where the alternative configuration is seen occasionally (72), but mostly not. So, we have nominally similar VLP and receptor-complex structures exhibiting different equilibrium states. Absent an understanding of how a finely balanced equilibrium is influenced, one should be cautious about attributing mechanistic significance, whether for packaging mutants or in receptor-binding. In summary, comparison of the Meyer *et al*. and Zhang *et al*. reconstructions reveals a putative conformational equilibrium in which N-terminally of Gly_236_, the structures diverge. Tracing towards the N-terminus, the chain either heads down βA towards the 5-fold pore, or to disordered structures near the 2-fold (Figure 4C-E). The volume of the partially ordered segment of the map is ∼12,000 Å³, corresponding to 89 typically sized amino acids. Thus, the disordered region, which is centered on a two-fold axis, could contain two copies of the N-terminal 35 residues of VP3 (before βB). Alternatively, each could contain a single copy of either VP2 (65 + 35 = 100 residues before the βA-βB turn) or part of VP1 (202 + 35 = 237 residues), noting that there are a total of ∼12 copies combined of VP1 and VP2, but 30 × 2-fold axes. For conformers headed towards the 5-fold, up to one in five would have access to the exterior through the pore, while others might be part of the “basket-like” disordered structure, seen surrounding the 5-fold on the inner surfaces of some, but not all AAVs (76).

### Comparison of single particle reconstructions for AAV5 bound with PKD1-2

A single particle analysis of the AAV5/PKD1-2 was also performed, using particles from the same grid that was subject to tomographic analysis. The reconstruction, at 2.88 Å resolution, agrees well with the previous 2.5 Å map (59), similarly resolving PKD1, but not PKD2. The new SPA and tomography data were collected from the same sample grid, so detection of PKD2 is a result of the technique not the sample (Figure 5B-D). While the high-resolution SPA structures of PKD1 are mostly very similar, there are differences in the N terminal residues (Figure 5A, arrows). This is at the same region where the two prior single particle analyses had been modeled differently (58, 59) in weak regions of their respective maps. The first β-strand of PKD1 is segmented, with βA followed by a break for a slight change of direction then βA’. The structures diverge upstream of the break near residue 315 (Fig. 5A, arrows). The N-terminal residue of the Silveria *et al*. construct, Val_311_, is not seen, but the rest of the structure is homologous to PKD PDBid 2yrl. The Zhang *et al*. construct starts upstream, so is longer at the N-terminus. It is modelled more like homolog PDBid 2y72, with βA occupying part of a different map feature, and connecting to βA’ with a non-β linker (residues 315 to 313). This configuration allows the authors to predict a contact near the 5-fold (AAVR Val_305_ & AAV5 Ser_319_), but the map at the N-terminus is weak and does not offer experimental support. At low contour in an unsharpened map, the new 2.88 Å SPA reveals additional extended chain / β-strand approximately parallel to βA’. The observation prompted retrospective examination of the prior 2.5 Å reconstruction (59). The same, hitherto unrecognized, feature looks very similar at low contour level in the unsharpened map. Furthermore, the map deposited by Zhang *et al*. was re-examined, and similar features were found (Figure 5C), only some of which are accounted by their different βA configuration. Given a lack of side chains, the identity of the extra features cannot be determined unambiguously. They are best described as a β-hairpin “U” (Figure 5B). Hypothetically, one could account for the two arms of the “U” separately with two additional configurations for βA-βA’ of AAVR PKD1. The melting of β-sheet hydrogen bonds to spring βA’ loose seems implausible and there is not the diminution of density for βA’ expected if βA’ had alternate conformers.

**Figure 5:**
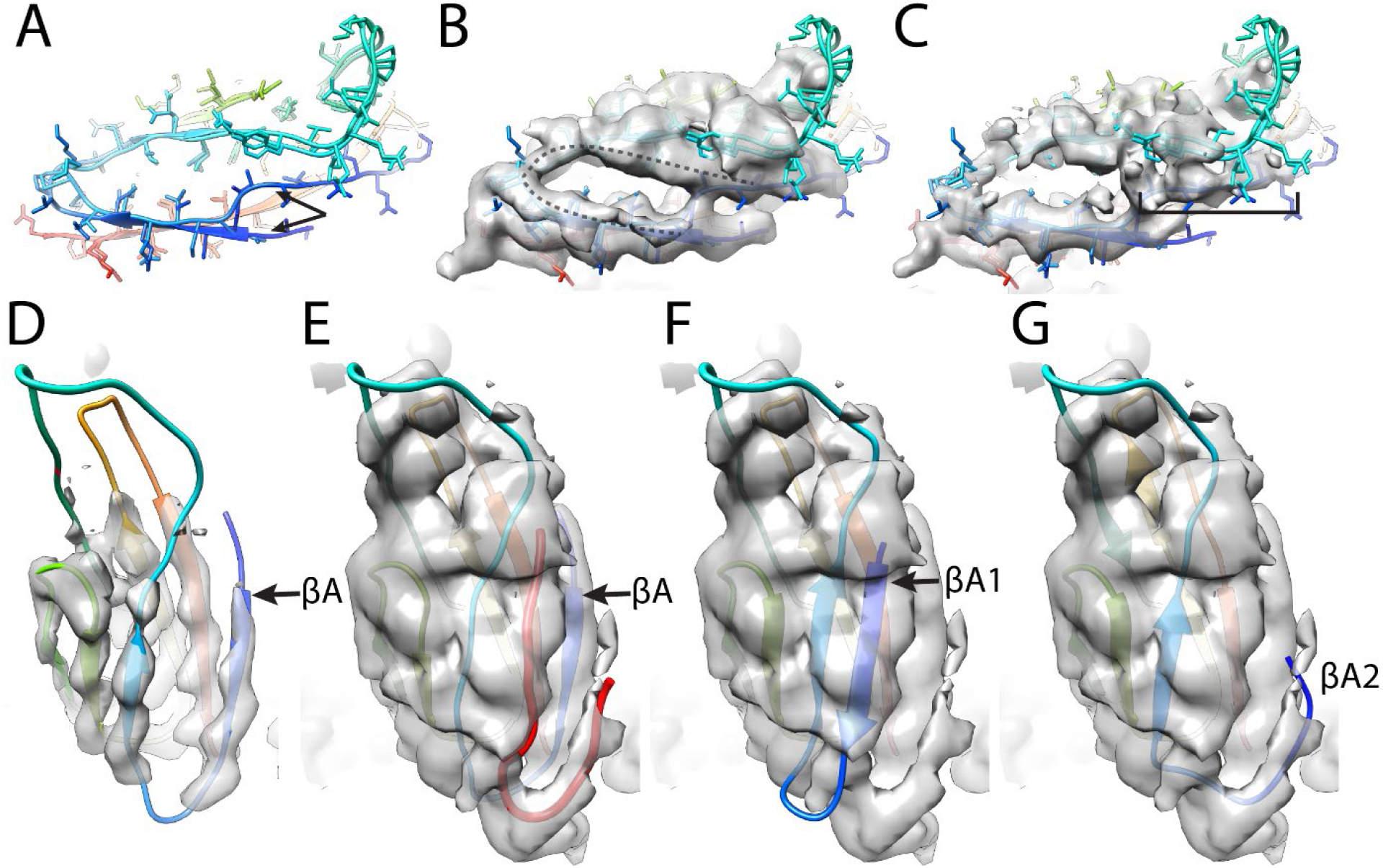
PKD1 structure from SPA of AAV5 complexes. A) Comparison of the Silveria et al. and Zhang et al. models of AAVR PKD1 as bound to AAV5. Arrows indicate the discrepancies between the N-terminal residues of the two models. B) Models in panel A compared to the cryo-EM SPA map determined here. Dashed line indicates unmodelled features at low contour. C) Models in panel A compared to the Zhang et al. map, the bracket indicating part of the dashed region modeled by Zhang et al., but not Silveria et al. D) At high contour, the current reconstruction supports the Silveria model as at least a dominant conformer. The first strand is labeled βA. E) Low contouring of the new map shows two putative β strands (red) continuing the β sheets on the top and bottom of the PKD1 structure. In this model βA is in the same position as in (D). This panel shows the most probable interpretation, but panels F & G present an alternative. F) Homologs in the PDB have structures similar to both Silveria and Zhang models and, less frequently, like PDBid 6aem. Overlay of this structure at 1.3 Å resolution shows how the longer of the unmodeled regions could be an alternative conformer of βA, labeled βA1. G) The shorter unmodeled region would have to be a 3^rd^ conformer of βA, labeled βA2.

Using high contour levels for the new map, we see that the predominant configuration of PKD1 is as modeled by Silveria *et al*. (Fig. 5). It is most likely that the new features belong to a single peptide distinct from that previously modeled. In the discussion, two possibilities will be considered, either that the unaccounted density is part of a second AAVR subunit, or that it is a fragment of hitherto unseen N-terminal regions of an AAV capsid protein.

## Discussion

Cryo-ET has allowed a subvolume classification that revealed the locations of receptor domains which were missing from the previous single particle analyses (SPA), even with attempted SPA subvolume classification (43, 77). For AAV5, PKD1 had been visualized by SPA (58, 59), but there had been no sign of PKD2. The cryo-ET showed PKD2 doubled back over the top of PKD1 in two orientations differing by ∼30°, likely neither of sufficient occupancy and order to be resolved by cryo-EM SPA. Such disorder and heterogeneity are consistent with PKD2 having few interactions, limited contacts with PKD1 near the interdomain hinge, and no contacts with AAV5 beyond the PKD1-2 domain linker for either of the classes. The distal locations of PKD2, revealed by cryo-ET, are also consistent with analysis of domain-deletion and chimeric domain-swapped mutants which indicated that PKD1, but not PKD2 have significant impact upon AAV5 cellular transduction (54).

AAV2 presented more of an enigma, because the same mutational analysis found that PKD2 was most important for AAV2 entry, but PKD1 also enhanced transduction, though to lesser extent. PKD2, the more critical for transduction, had previously been resolved by SPA (43, 57), but not the “accessory” PKD1. Prior to the SPA structures of AAV5-AAVR complexes (58, 59), we hypothesized that the unseen PKD1 might be interacting loosely with AAV2 at a site corresponding to the (yet to be determined) AAV5-PKD1 interface. The cryo-ET shows that none of PKD1 locations of any of the four classes in the AAV2 complex bear any resemblance to PKD1 as bound by AAV5.

However, one of the four PKD1 classes has some direct contact with AAV2 proteins. This is consistent with PKD1 playing an accessory role not strictly required for, but enhancing cellular transduction. Note, however, that only one of the four AAV2 classes appears to make contact and the contact is not extensive. Thus, it is not surprising that there can be the observed wide-ranging heterogeneity in domain orientation, the four classes spanning a 120° rotation about the interdomain hinge. While we would expect the more populated orientations to rise to the top of classification, there might well be diversity beyond the four discretely classed orientations (as indicated by the XL-MS), and it is not surprising that PKD1 was not detectable by SPA. Clearly the level of interactions between PKD1 and either AAV2 or PKD2 are insufficient to restrict conformational heterogeneity, so one wonders whether the interactions with AAV2 can be strong enough to have a measurable direct impact upon transduction through avidity. It seems more likely that either PKD1 increases the availability or stability of AAVR in a state compatible with the binding of AAV2 to PKD1, or that there is a different step in AAV entry in which PKD1 has a role.

Completely unanticipated was the unmodeled density on the inside surface of the AAV2/PKD1-2 complex (but not the AAV5/PKD12 complex). It correlates inversely with the strength of βA density, density for βA being much weaker when the unmodeled features are seen. Thus, it appears that we are observing an equilibrium between two states, one with an ordered βA extending from the 5-fold region, and the other with a partially ordered N-terminal region coming from the inner surface protrusion, skipping βA, and joining the jellyroll fold capsid protein at the βA-βB hairpin turn. The volume of the inner protrusion is commensurate with that expected of the N-terminal 35 residues of two VP3 meeting at a 2-fold axis, although one cannot rule out partial occupancy by VP2 or VP1. Whether and how this equilibrium in N-terminus location is influenced by receptor-binding far away on the outside surface are unknown.

Another surprise was the previously unseen fragments of β-strand structure adjacent to PKD1 in its complex with AAV5. They lacked distinctive features to identify by sequence. Nevertheless, there are a limited number of plausible possibilities. The N-terminal regions of the capsid proteins have never been seen at high resolution. While in this study partially ordered structures were seen on the interior surface of AAV2, crystal structures of some AAVs and autonomous parvoviruses have indicated that a fraction of N-termini (of at least VP3) might be external: partially ordered density running down the 5-fold pore from the outside is interpreted as the connection to the start of the β-barrel on the inside surface (78-81). The absence of density on the 5-fold axis in the AAV5 single particle analyses lessens the likelihood that the unaccounted features are previously unresolved N-terminal parts of the viral protein outside the capsid.

Alternatively, the extra peptides could come from unmodeled regions of AAVR. Dimers and higher oligomers are seen in preparations of PKD1-2 constructs (and MBP-PKD1-5 fusions) (43, 61). To date, AAVR dimers have not been observed bound to AAV5, but one cannot exclude the possibility that a small fraction of receptors in the complex are dimerized, with disorder that precludes EM observation of most of the second subunit.

This work is a testament to the value of combining multi-technique, multi-scale approaches for flexible complexes, and in recognizing gaps in our understanding through exclusive reliance on high resolution structure. A plan for multiple contingencies involved not only integration of different EM techniques, but also upstream redundancy in expression constructs, both of which were needed for a more robust and holistic understanding. It is noted that the first application of cryo-ET, to a complex of AAV2 with a PKD1-5 MBP fusion construct, led to a very low resolution visualization that lacked domain definition or perception of conformational heterogeneity (43). It was only with a smaller construct, His_6_-PKD1-2, that higher binding occupancy was achieved and conformational heterogeneity from domains 3-5 was eliminated, making it possible to classify the remaining heterogeneity and resolve distinct configurations for the two proximal domains. On the technical side, it is noted that fully automated classification of subvolume tomograms within a symmetrical particle was not yet possible. It is hoped that examples like this will inspire on-going algorithm development, so that future applications will not be limited by the laboriousness of interactive classification.

## Acknowledgements

Funding for this research was provided by:

National Institutes of Health: R35GM122564 (M.S.C.), and in part by R01GM108753 (S.M.S.). Maps for the subvolume averages and single particle reconstruction will be made available at the EM data resource database.

## Author contributions

Conceptualization and Methodology: S.M.S. and M.S.C.; Funding acquisition: M.S.C.; Sample preparation: M.A.S.; Data collection: G.H.; Data analysis: G.H. and M.A.S.; Writing – original draft preparation: G.H., S.M.S. and M.S.C.; Writing – review and editing: all authors.

